# Computationally Driven Design of Novel BACE-1 Inhibitors for Alzheimer’s Disease

**DOI:** 10.1101/2023.11.15.567308

**Authors:** Megan M. Chinn, Diego Lopez Mateos, Mary A. Riley, Justin B. Siegel

## Abstract

Alzheimer’s disease is the most common form of dementia affecting 35 million people globally. One of the major efforts in the development of a treatment for Alzheimer’s is to reduce the rate of plaque formation, the common hallmark of Alzheimer’s disease. The protease BACE-1 has been demonstrated to play a role in catalyzing plaque formation and is therefore a major drug target. Here we report potential new drug candidates building upon Verubecestat, developed through computationally driven molecular modeling methods. Both designed molecules have improved docking scores relative to Verubecestat when modeled in the BACE-1 active site, and therefore present potential new leads for more effective therapeutics to combat Alzheimer’s disease.

## INTRODUCTION

Alzheimer’s disease is a neurodegenerative and polygenic disease. It is the most common form of dementia, and patients diagnosed with it experience continuous cognitive deterioration leading to memory loss and it can potentially be fatal. Worldwide, about 35 million people suffer from Alzheimer’s disease, and by 2030, it’s estimated that 74.7 million people will be diagnosed.^1^

Alzheimer’s disease is caused by various factors, including the presence of extracellular amyloid plaques as well as neuroinflammation and loss of neurotransmitters in the brain.^1^ These Aβ (amyloid-β) peptides are insoluble, neurotoxic proteins that cause plaque deposition and neurodegeneration.^2^ The Aβ peptides are produced by the proteolytic cleavage of the amyloid precursor protein (APP) by the enzyme beta-secretase 1 (BACE-1). The inhibition of BACE-1 by blood-brain permeable inhibitors, can prevent Aβ production with the potential to improve cognitive and behavioral deficiencies.^3^

BACE-1 is an aspartyl protease type 1 transmembrane protein of the pepsin family. It’s present in the Golgi apparatus, late endosomes, pancreas, but most importantly in the brain, particularly in various neuronal cell types. It catalyzes the cleavage of APP; and a mutation in the substrate APP increases BACE-1’s proteolytic cleavage of it, resulting in an increased production of Aβ peptides and the early onset of Alzheimer’s.^4^

A notable mention is BACE-2, a homolog to BACE-1. It is present in peripheral tissues and the brain, but mainly in the kidneys. It cleaves the same APP substrate and produces Aβ peptides as well, however, because it is less abundant in the brain than BACE-1, its generation of Aβ peptides is less noticeable. This suggests that inhibiting drugs should mainly target BACE-1 to efficiently combat Alzheimer’s disease.^5^

With the large number of people affected by this disease every year, it is no surprise that there have been multiple attempts at creating drugs to cure Alzheimer’s. Figure 1 includes four examples of drugs on the market for this disease including Atabecestat, Lanabecestat, CTS-21166, and Verubecestat.

**Figure 1.**
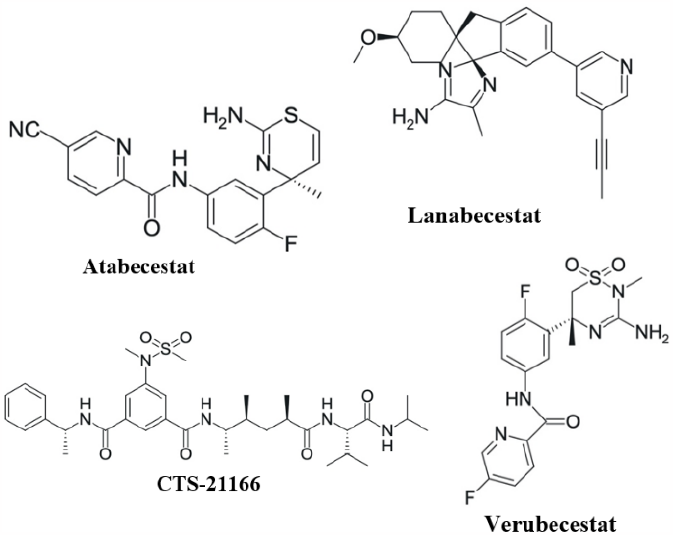
The 2D structures of Alzheimer’s drugs Atabecestat, Lanabecestat, CTS-21166, and Verubecestat.

All four molecules inhibit BACE-1, but also have many off-target effects. One hypothesis is that this is due to the lack of specificity, as Lanabecestat, Atabecestat, and Verubecestat can also inhibit BACE-2.^6-8^ Lanabecestat also has an off-target effect on hypothalamic leptin sensitivity (affecting body weight) due to lowered BACE-1 activity.^6^ Atabecestat causes hepatotoxicity, increasing liver enzyme levels and affecting liver function.^7^ Verubecestat and Atabecestat also inhibit the hERG channel, leading to QT_C_ prolongation and cardiac arrhythmias (irregular heart-beat).^8,9^ CTS-21166 is found to alter myelination in peripheral nerves.^10^ These off-target effects leave much to be desired. However, the research on these drugs opens the door to improvements that can be made to create other potential candidates.

In this study, we propose two novel BACE-1 inhibitor drug candidates based on the known drug Verubecestat. The first was created with the program vBrood, and the second based on chemical intuition. Their absorption, distribution, metabolism, excretion, and toxicity (ADMET) characteristics were analyzed. Both were docked into the BACE-1 active site and found to bind better than the original drug candidate and therefore are promising for improved drugs to combat Alzheimer’s disease.

## METHODS

For the control experiment, RCSB Protein Data Bank was accessed for the crystal structure of BACE-1 (PDB: 7D2V).^11,12^ PyMOL was used to analyze the crystal structure active site and measure the distances of ligand-to-residue interactions.^13^

The baseline drugs and novel candidates were built and optimized with Gaussian 09W^14^ and Gaussview.^15^ Open-Eye-OMEGA^16^ was then used to generate the conformer libraries. While OpenEye-FILTER^16^ software was used to generate the ADMET properties and run the Lipinski’s rule of 5 test on all potential drugs. VIDA^17^ allowed for visualization and analyzation of these drugs.

For docking, Fred Make Receptor^18^ allowed for designation of the binding site in the PDB structure. Then Fred was used again to dock all drug molecules into this binding site and generate a docking report.

To generate Candidate 1, vBrood^19^ was used with the baseline Verubecestat drug to create a library of potential novel drugs with bioisosteric replacements, from which one was chosen as the candidate. Candidate 2 was designed and built via chemical intuition. Both candidates were then docked and analyzed the same way as the baseline drugs. SciFinder^20^ was used to confirm that Candidates 1 and 2 had not been proposed before.

## RESULTS AND DISCUSSION

Four pre-existing drug inhibitors of BACE-1 were studied and analyzed in preparation for generating novel drug Candidates 1 and 2. First, docking reports were generated for Verubecestat, Lanabecestat, Atabecestat, and CTS-21166 (Table 1).

**Table 1.** Docking Report of the Four Drug Candidates.

| Name | Docking Score | Shape | Hydrogen Bonding |
| --- | --- | --- | --- |
| Verubecestat | -8.41 | -11.34 | -1.47 |
| Lanabecestat | -9.47 | -12.39 | -2.87 |
| Atabecestat | -8.71 | -8.32 | -6.35 |
| CTS-21166 | -2.43 | -8.49 | -2.55 |

From the above table, Lanabecestat had the best (most negative) docking score of -9.47, indicating it had the best predicted interactions and fit with the binding site. Lanabecestat also had the best shape score of -12.39, which probably contributed to its low docking score. Atabecestat had the second-best docking score at -8.71 and also had the best hydrogen bond score of -6.35, suggesting it can form more hydrogen bonds within the active site. With a docking score very close to Atabecestat, Verubecestat is at -8.41, but somehow is predicted to form the worst hydrogen bonds with a score of -1.47. On the other hand, CTS-21166 was predicted to bind the worst with the binding site, indicative of its unfavorable docking and hydrogen bonding scores of -2.43 and -2.55, respectively.

The drugs were further analyzed for features that would affect their potential ADMET behavior (Table 2), where CTS-21166 was the only drug that failed to meet Lipinski’s rules, as it had three violations including molecular weight over 500 g/mol, XLogP over 5, and number of hydrogen bond acceptors over 10. Fortunately, the three other drugs passed with no Lipinski violations, and thus increasing the probability of being orally administered.

**Table 2.** ADMET Properties of the Four Drug Candidates.

| Name | Molecular Weight | Lipinski H Bond Acceptor | Lipinski H Bond Donor | XLogP | PSA |
| --- | --- | --- | --- | --- | --- |
| Verubecestat | 409.41 | 8 | 2 | -0.04 | 117.75 |
| Lanabecestat | 413.53 | 5 | 2 | 4.04 | 74.47 |
| Atabecestat | 368.41 | 6 | 3 | 1.84 | 105.77 |
| CTS-21166 | 656.86 | 11 | 4 | 6.25 | 153.78 |

Since Verubecestat had no Lipinski violations and the lowest hydrogen bonding score, it was hypothesized that increasing this score would improve the docking score, while remaining orally bioavailable with the addition of new molecular features. Specifically, with just two hydrogen bond donors and a molecular weight of 409.41 g/mol, Verubecestat had the capacity to be structurally modified to increase the number of donating groups with a low chance of surpassing the Lipinski molecular weight of 500 g/mol.

A previously solved crystal structure with Verubecestat bound in the active site (PDB: 7D2V)^11,12^ was analyzed in the active site using PyMOL to visualize all interactions (Figure 2). As shown, the two fluorine atoms both formed hydrogen bonds with surrounding residues, THR 232 and TYR 71, at the lengths of 3.4Å and 3.8Å, respectively.

**Figure 2.**
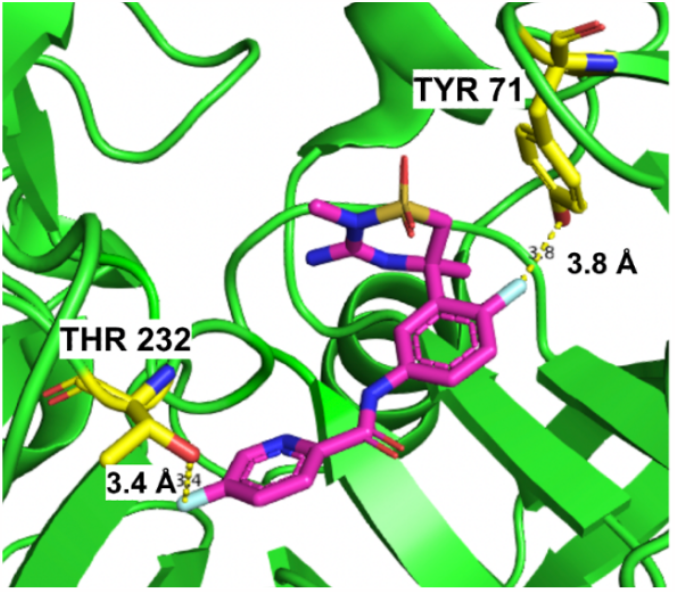
Verubecestat (pink) interactions with THR 232 and TYR 71 (yellow) in the BACE-1 active site.

### Candidate 1: Computationally Driven Design

Using the program vBrood, various bioisosteric replacements were made on Verubecestat to generate a list of potential candidates. They were analyzed in VIDA and docked in the BACE-1 active site using FRED. The docking scores were generated and the best score of a Verubecestat bioisosteric replacement was found. As a side note, this drug was also run through SciFinder to ensure it wasn’t proposed before.

The new Candidate 1 (Figure 3A) contained the original Verubecestat structure with a replacement of the sulfur-containing heterocycle with a ring containing two nitrogen atoms and an additional cyclopropane. Table 3 shows the resulting docking score of -12.70, a 51% improvement over the Verubecestat score of -8.41. There was an unfavorable increase in shape score (-11.34 to -8.39), suggesting it didn’t bind as well as Verubecestat. Though, this is compensated by a dramatically favorable decrease in hydrogen bonding score, changing from -1.47 to -11.40.

**Table 3.**
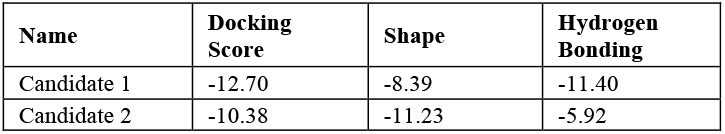
Docking Report of Candidates 1 & 2.

| Name | Docking Score | Shape | Hydrogen Bonding |
| --- | --- | --- | --- |
| Candidate 1 | -12.70 | -8.39 | -11.40 |
| Candidate 2 | -10.38 | -11.23 | -5.92 |

**Figure 3.**
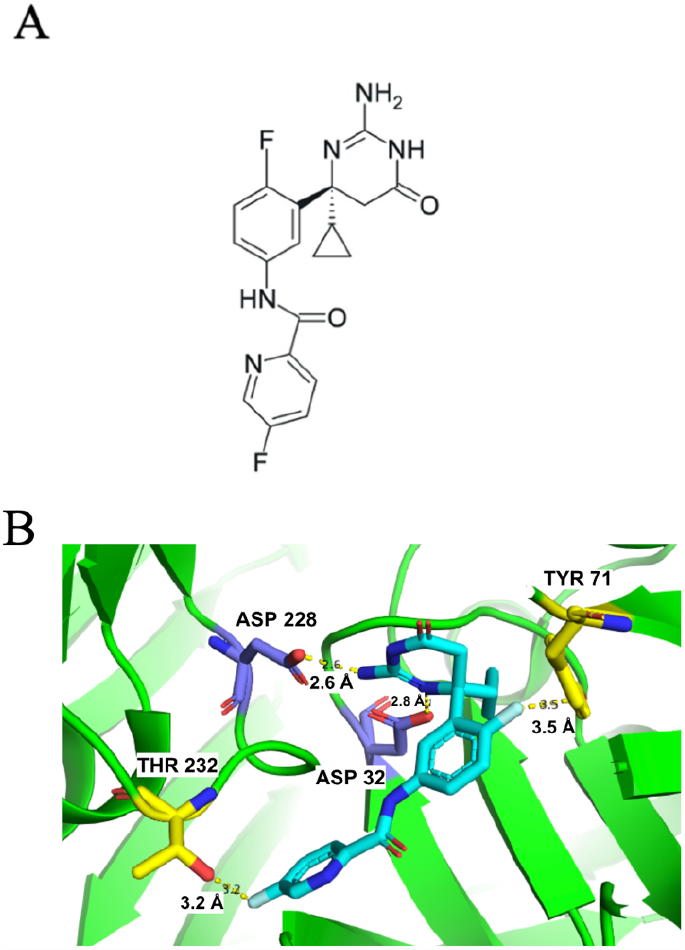
**A)** The structure of Candidate 1. **B)** Candidate 1 (blue) existing interactions with THR 232 and TYR 71 (yellow) and new interactions with ASP 228 and ASP 32 (purple) in the BACE-1 active site.

This docking score improvement can be explained using PyMOL to visualize Candidate 1 interactions with surrounding residues (Figure 3B). Candidate 1 had the same existing interactions the original Verubecestat had, however, the bond lengths were shorter, strengthening the hydrogen bonds formed. Specifically, these hydrogen bonds shortened from 3.8Å to 3.5Å for TYR 71 and from 3.4Å to 3.2Å for THR 232. It’s probable that these shorter bond lengths partially contributed to the decrease in hydrogen bonding for Candidate 1. It was also observed that Candidate 1 formed two new hydrogen bonds with ASP 228 and ASP 32 at bond lengths of 2.6Å and 2.8Å, respectively.

Shown in Table 4 are the ADMET properties of Candidate 1, which are important to consider when comparing the viability of this drug to Verubecestat. The molecular weight of Candidate 1 was lower due to the removal of sulfur and oxygen, 386.38 g/mol versus the original 409.41 g/mol. There were a higher number of total hydrogen bond acceptors and donors (8 HBA and 3 HBD) compared to Verubecestat (8 HBA and 2 HBD) from the additional amine groups. Candidate 1 had a higher XLogP value of 0.99 and a lower polar surface area (PSA) value of 111.08, while Verubecestat had respective values of -0.04 and 117.75. This increase in XLogP could be due to the removal of the SO_2_ group, allowing access to the surrounding carbons in the ring. The decrease in PSA could be due to the removal of the oxygen group. Overall, Candidate 1 meets all Lipinski’s rules, while displaying improved docking scores, interactions, and ADMET properties over the original drug.

**Table 4.** ADMET Properties of Candidates 1 & 2.

| Name | Molecular Weight | Lipinski H Bond Acceptors | Lipinski H Bond Donors | XLogP | PSA |
| --- | --- | --- | --- | --- | --- |
| Candidate 1 | 386.38 | 8 | 3 | 0.99 | 111.08 |
| Candidate 2 | 390.34 | 8 | 3 | 0.65 | 111.63 |

### Candidate 2: Chemical Intuition Driven Design

Candidate 2 (Figure 4A) was intuitively designed around Verubecestat, with inspiration from Candidate 1. Both Verubecestat and Candidate 1 had interactions with THR 232 and TYR 71, therefore, it was hypothesized that Candidate 2 should maintain a similar structure of two fluorine-containing rings conjugated by a secondary amine and ketone. Thus, the sulfur-nitrogen heterocycle of Verubecestat was again the main target for modification by adding a hydroxyl and methyl group, hopefully lowering the docking score. Oxygens replaced the primary and tertiary amines in an attempt to develop a more electronegative nature with stronger hydrogen bond potential. The final modification was to replace the sulfate group with a primary amine for further increased hydrogen bonding potential. Candidate 2 was run through SciFinder to confirm it had not been previously developed.

**Figure 4.**
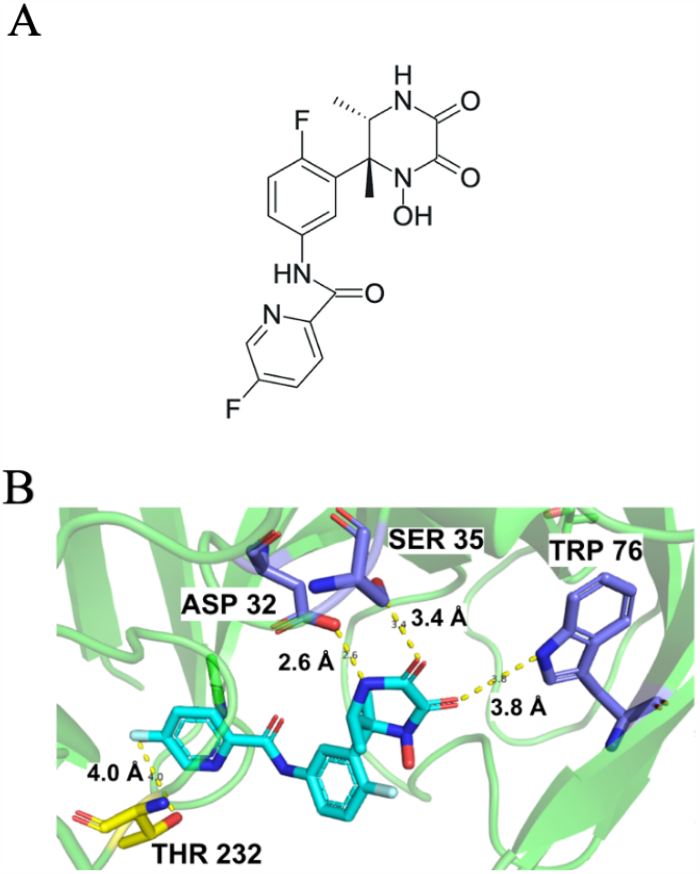
**A)** Structure of Candidate 2. **B)** Candidate 2 (blue) existing interaction with THR 232 (yellow) and new interactions with ASP 32, SER 35, and TRP 76 (purple) in the BACE-1 active site.

The docking score for Candidate 2 is shown in Table 3 with a 23% improvement over Verubecestat, decreasing from -8.41 to -10.38. The hydrogen bonding score also decreased significantly from -1.47 to -5.92, while the shape score increased slightly from -11.34 to -11.23.

The improvement in docking score can be visualized via molecular interactions using PyMOL (Figure 4B). Candidate 2 had the same existing interaction with THR 232 that Verubecestat had, but the distance increased to 4.0Å, thus, turning from a hydrogen bond into an electrostatic interaction. Similarly, the original Verubecestat interaction between the second fluorine and TYR 71, increased to a distance too far for any interactions to occur (not shown). This loss of an interaction can be justified by the ring modifications that altered the orientation of Candidate 2 binding.

Fortunately, new interactions include several hydrogen bonds, one with TRP 76 at a distance of 3.8Å, another with SER 35 at a shorter distance of 3.4Å, and lastly an even stronger hydrogen bond with ASP 32 at a distance of 2.6Å. So, although Candidate 2 lost two hydrogen bond interactions, it gained three moderately strong hydrogen bonds which all contributed to the decrease in hydrogen bonding score to -5.92.

The ADMET properties for Candidate 2 were generated and are shown in Table 4. The molecular weight decreased from 409.41 g/mol to 390.34 g/mol due to the replacements. While the total number of hydrogen bond donors and acceptors increased from 10 to 11 due to the addition of the hydroxyl group. The XLogP increased from -0.04 to 0.65, possibly due to the removal of the sulfate which allows for access to the carbons in the ring. While the PSA decreased from 117.75 to 111.633, which could be the result of removing an amine group. However, despite these differences, Candidate 2 has potential to retain its oral bioavailability as it doesn’t violate any of Lipinski’s rules and is thus a viable potential drug candidate for Alzheimer’s disease.

## CONCLUSION

With 35 million people currently experiencing the effects of Alzheimer’s disease, and many more projected in the coming years, it makes sense that many studies have been conducted to combat it.^1^ To help contribute to finding a cure, two novel drugs were designed, one using computational methods and the other with chemical intuition.

The computationally driven designed drug, Candidate 1, showed the most promise with a 51% favorable decrease in docking score (-8.41 to -12.70). It also had a vast improvement in hydrogen bonding score (-1.47 to -11.40). Furthermore, Candidate 1 had no Lipinski violations, thus, it is an exciting lead future Alzheimer’s treatment.

Similarly, Candidate 2 had a 23% improvement in docking score (-8.41 to -10.38) as well as a decrease in hydrogen bonding (-1.47 to -5.92). It had no Lipinski violations, and although it had a lower docking score than Candidate 1, Candidate 2 was still an improvement on Verubecestat and is therefore a potential lead for future treatments. In conclusion, Candidates 1 and 2 are two promising leads for further research towards better therapeutic options for Alzheimer’s patients.

## AUTHOR INFORMATION

### Present Addresses

† Department of Chemistry, University of California Davis, Davis, California, United States of America, Department of Biochemistry and Molecular Medicine, University of California, Davis, Davis, California, United States of America, Genome Center, University of California Davis, Davis, California, United States of America.

### Author Contributions

Research was designed by all authors; all experiments were carried out by M.C. The manuscript was written through contributions of all authors. All authors have given approval to the final version of the manuscript.

## ACKNOWLEDGMENT

Research reported in this publication was supported by UC Davis. This study was derived from a course based undergraduate research study conducted in Chemistry 130B at UC Davis.

## Notes

### Competing Interest Statement

The authors have declared no competing interest.

## REFERENCES

1. Das, B.; Yan, R. A Close Look at BACE1 Inhibitors for Alzheimer’s Disease Treatment. CNS Drugs 2019, 33 (3), 251–263.

2. Moussa-Pacha, N. M., Abdin, S. M., Omar, H. A., Alniss, H., & Al-Tel, T. H. BACE1 inhibitors: Current status and future directions in treating alzheimer’s disease. Medicinal Research Reviews. 2019, 40(1), 339–384. 10.1002/med.21622

3. Fujimoto, K.; Yoshida, S.; Tadano, G.; Asada, N.; Fuchino, K.; Suzuki, S.; Matsuoka, E.; Yamamoto, T.; Yamamoto, S.; Ando, S.; Kanegawa, N.; Tonomura, Y.; Ito, H.; Moechars, D.; Rombouts, F.; Gijsen, H.; Kusakabe, K. Structure-Based Approaches to Improving Selectivity through Utilizing Explicit Water Molecules: Discovery of Selective β-Secretase (BACE1) Inhibitors over BACE2. J Med Chem. 2021, 64 (6).

4. Luo, X.; Yan, R. Inhibition of BACE1 for therapeutic use in alzheimer’s disease. International journal of clinical and experimental pathology. 2010, 3 (6).

5. Ahmed, R. R.; Holler, C. J.; Webb, R. L.; Li, F.; Beckett, T. L.; Murphy, M. P. BACE1 And BACE2 Enzymatic Activities in Alzheimer’s Disease. Journal of Neurochemistry 2010, 112 (4), 1045–1053.

6. Wessels, A. M.; Tariot, P. N.; Zimmer, J. A.; Selzler, K. J.; Bragg, S. M.; Andersen, S. W.; Landry, J.; Krull, J. H.; Downing, A. C. M.; Willis, B. A.; Shcherbinin, S.; Mullen, J.; Barker, P.; Schumi, J.; Shering, C.; Matthews, B. R.; Stern, R. A.; Vellas, B.; Cohen, S.; MacSweeney, E.; Boada, M.; Sims, J. R. Efficacy and Safety of Lanabecestat for Treatment of Early and Mild Alzheimer Disease. JAMA Neurology 2020, 77 (2), 199.

7. Koriyama, Y.; Hori, A.; Ito, H.; Yonezawa, S.; Baba, Y.; Tanimoto, N.; Ueno, T.; Yamamoto, S.; Yamamoto, T.; Asada, N.; Morimoto, K.; Einaru, S.; Sakai, K.; Kanazu, T.; Matsuda, A.; Yamaguchi, Y.; Oguma, T.; Timmers, M.; Tritsmans, L.; Kusakabe, K.; Kato, A.; Sakaguchi, G. Discovery of Atabecestat (JNJ-54861911): A Thiazine-Based β-Amyloid Precursor Protein Cleaving Enzyme 1 Inhibitor Advanced to the Phase 2b/3 EARLY Clinical Trial. Journal of Medicinal Chemistry. 2021, 64 (4).

8. Forman, M.; Palcza, J.; Tseng, J.; Stone, J. A.; Walker, B.; Swearingen, D.; Troyer, M. D.; Dockendorf, M. F. Safety, Tolerability, and Pharmacokinetics of the β-Site Amyloid Precursor Protein-Cleaving Enzyme 1 Inhibitor Verubecestat (Mk -8931) in Healthy Elderly Male and Female Subjects. Clinical and Translational Science 2019, 12 (5), 545–555.

9. Hsiao, C.-C.; Gijsen, H. Atabecestat. https://www.sci-encedirect.com/topics/pharmacology-toxicology-and-phar-maceutical-science/atabecestat

10. Yan, R. Stepping Closer to Treating Alzheimer’s Disease Patients with BACE1 Inhibitor Drugs. Translational Neurodegeneration 2016, 5 (1).

11. H.M. Berman, J. Westbrook, Z. Feng, G. Gilliland, T.N. Bhat, H. Weissig, I.N. Shindyalov, P.E. Bourne. The Protein Data Bank. Nucleic Acids Research [Online], 235–242. https://www.rcsb.org/ (accessed March 8, 2023).

12. Fujimoto, K., Yoshida, S., Tadano, G., Asada, N., Fuchino, K., Suzuki, S., Matsuko N., Yamamoto, T., Yamamoto, S., Ando, S., Kanegawa, N., Tonomura, Y., Ito, H., Moechars, D., Rombouts, F.J.R., Gijsen, H.J.M., Kusakabe, K.I. (2020) Crystal Structure of BACE1 in complex with N-{3-[(5R)-3-amino-2,5-dimethyl-1,1-dioxo-5,6-dihydro-2H-1lambda6,2,4-thiadiazin-5-yl]-4-fluorophenyl}-5-fluoro-pyridine-2-carboxamide. doi: 10.2210/pdb7D2V/pdb

13. The PyMOL Molecular Graphics System, Version 2.0 Schrödinger, LLC.

14. Gaussian 09, Revision W, M. J. Frisch, G. W. Trucks, H. B. Schlegel, G. E. Scuseria, M. A. Robb, J. R. Cheeseman, G. Scalmani, V. Barone, G. A. Petersson, H. Nakatsuji, X. Li, M. Caricato, A. Marenich, J. Bloino, B. G. Janesko, R. Gomperts, B. Mennucci, H. P. Hratchian, J. V. Ortiz, A. F. Izmaylov, J. L. Sonnenberg, D. Williams-Young, F. Ding, F. Lipparini, F. Egidi, J. Goings, B. Peng, A. Petrone, T. Henderson, D. Ranasinghe, V. G. Zakrzewski, J. Gao, N. Rega, G. Zheng, W. Liang, M. Hada, M. Ehara, K. Toyota,R. Fukuda, J. Hasegawa, M. Ishida, T. Nakajima, Y. Honda, O. Kitao, H. Nakai, T. Vreven, K. Throssell, J. A. Montgomery, Jr., J. E. Peralta, F. Ogliaro, M. Bearpark, J. J. Heyd, E. Brothers, K. N. Kudin, V. N. Staroverov, T. Keith, R. Kobayashi, J. Normand, K. Raghavachari, A. Rendell, J. C. Burant, S. S. Iyengar, J. Tomasi, M. Cossi, J. M. Millam, M. Klene, C. Adamo, R. Cammi, J. W. Ochterski, R. L. Martin, K. Morokuma, O. Farkas, J. B. Foresman, and D. J. Fox, Gaussian, Inc., Wallingford CT, 2016.

15. Nielsen, A.B. and Holder, A.J. (2009) Gauss View 5.0, User’s Reference. GAUSSIAN Inc., Pittsburgh

16. ROCS 3.5.1.2: OpenEye Scientific Software, Santa Fe, NM. http://www.eyesopen.com.

17. VIDA, Version 4.4.0.4; OpenEye Scientific Software: Santa Fe, NM, 2018.

18. McGann, M. FRED Pose Prediction and Virtual Screening Accuracy. J. Chem. Inf. Model., 2011, 51, 578–596. DOI: 10.1021/ci100436p

19. vBrood, Version 3.1.1.2; OpenEye Scientific Software: San-ta Fe, NM.

20. SciFinder; Chemical Abstracts Service: Columbus, OH; carbon-13 NMR spectrum; spectrum ID CC-03-C_SPC-3734; RN 50-52-2; https://scifinder.cas.org (accessed March 15, 2023).

